# The combination of Bromelain and Acetylcysteine (BromAc) synergistically inactivates SARS-CoV-2

**DOI:** 10.1101/2020.09.07.286906

**Authors:** Javed Akhter, Grégory Quéromès, Krishna Pillai, Vahan Kepenekian, Samina Badar, Ahmed H. Mekkawy, Emilie Frobert, Sarah J Valle, David L. Morris

**Affiliations:** Department of Surgery, St George Hospital, Sydney NSW Australia; Mucpharm Pty Ltd, Australia; CIRI, Centre International de Recherche en Infectiologie, Team VirPatH, Univ Lyon, Inserm, U1111, Université Claude Bernard Lyon 1, CNRS, UMR5308, ENS de Lyon, F-69007, Lyon, FR; Hospices Civils de Lyon, Lyon, France; EMR 3738 (CICLY), Lyon 1 Université, Lyon, France; Laboratoire de Virologie, Institut des Agents Infectieux (IAI), Hospices Civils de Lyon, Groupement Hospitalier Nord, F-69004, Lyon, FR; University of New South Wales, St George & Sutherland Clinical School, Sydney NSW Australia

**Keywords:** SARS-CoV-2, Bromelain, Acetylcysteine, BromAc, drug repurposing

## Abstract

**Background and objectives:** SARS-CoV-2 infection is the cause of a worldwide pandemic, currently with limited therapeutic options. Whilst vaccines are at the forefront of the therapeutic initiative, drug repurposing remains a promising approach for SARS-CoV-2 treatment. BromAc (Bromelain & Acetylcysteine) has synergistic action against glycoproteins by the synchronous breakage of glycosidic linkages and disulfide bonds. The spike protein of SARS-CoV-2, formed of glycoprotein and disulfide bridges for stabilization, represents an attractive target as it is essential for binding to the ACE2 receptor in host cells present in nasal mucosa. We sought to determine the effect of BromAc on the Spike and Envelope proteins and its potential to reduce infectivity in host cells.

**Design:** Recombinant Spike and Envelope proteins were treated by single agent and combination BromAc at 50 and 100 µg/20mg/mL and analyzed by electrophoresis. Ultraviolet analysis of disulfide bond reduction was performed for both Spike and Envelope proteins after treatment with Acetylcysteine. *In vitro* whole virus culture inactivation of pre-treated wild type and an S1/S2 Spike mutant SARS-CoV-2 with BromAc from 25 to 250 µg/20mg/mL was measured by cytopathic effect, cell lysis assay, and replication capacity by RT-PCR.

**Results:** Recombinant Spike and Envelope SARS-CoV-2 proteins were fragmented by BromAc at both 50 and 100 µg/20mg/mL whilst single agents had minimal effect. Spike and Envelope protein disulfide bonds were reduced by Acetylcysteine. *In vitro* whole virus culture of both wild type and Spike mutant SARS-CoV-2 demonstrated a concentration-dependent inactivation from BromAc treatment but not from single agents.

**Conclusion:** BromAc disintegrates SARS-CoV-2 Spike and Envelope proteins. *In vitro* tests on whole virus support this finding with inactivation of its replication capacity most strongly at 100 and 250 µg/20mg/mL BromAc, even in Spike mutant virus. Clinical testing through nasal administration in patients with early SARS-CoV-2 infection is imminent.

**Author Summary:** There is currently no suitable therapeutic treatment for early SARS-CoV-2 aimed to prevent disease progression. BromAc is under clinical development by the authors for mucinous cancers due to its ability to alter complex glycoproteins structure. The potential of BromAc on SARS-CoV-2 Spike and Envelope glycoproteins stabilized by disulfide bonds was examined and found to disintegrate recombinant Spike and Envelope proteins whilst reducing disulfide stabilizer bridges. BromAc also showed an inhibitory effect on wild-type and Spike mutant SARS-CoV-2 by inactivation of its replication capacity *in vitro*. Hence, BromAc may be an effective therapeutic agent for early SARS-CoV-2 infection, despite mutations, and even have potential as a prophylactic in people at high risk of infection.

## Introduction

The recently emergent SARS-CoV-2 (severe acute respiratory syndrome coronavirus 2), also known as COVID-19, ranges from asymptomatic to severe lethal forms with a systemic inflammatory response syndrome. As of December 1^st^, 2020, over 63 million people worldwide have been infected, with an estimated overall mortality of 2.3% (1). There are currently few therapeutic agents proven to be beneficial in reducing early and late stage disease progression (2). While there are fortunately many vaccine candidates, the widespread availability for vaccination may not be immediate, the length of immune protection may be limited (3, 4) and efficacy of the vaccine may be reduced by viral mutations. The continued exploration of effective treatments is still needed.

Structurally, SARS-CoV-2 contains surface Spike proteins, Membrane proteins, and Envelope proteins, as well as internal Nucleocapsid proteins that contain the RNA. The Spike protein is a homotrimer glycoprotein complex with different roles accomplished through dynamic conformational modifications, based in part on disulfide bonds (5). It allows the infection of target cells by binding to the human angiotensin-converting enzyme (ACE2) receptors, among others, which triggers proteolysis by transmembrane protease serine 2 (TMPRSS2), furin, and perhaps other proteases, leading to capsid disintegration (6, 7).

Bromelain is extracted mainly from the stem of the pineapple plant (*ananas comosus*) and contains a number of enzymes that gives it the ability to hydrolyze glycosidic bonds in complex carbohydrates (8). As a therapeutic molecule, it is used for debriding burns. Acetylcysteine is a powerful antioxidant, commonly nebulized into the airway for mucus accumulation, and as a hepatoprotective agent in paracetamol overdose. Most importantly, Acetylcysteine reduces disulfide bonds (9). The combination of Bromelain with Acetylcysteine (BromAc) exhibits a synergistic mucolytic effect to treat mucinous tumors (10, 11) and as a chemosensitizer of several anticancer drugs (12). These different actions are due to the ability of BromAc to disintegrate molecular structures of complex glycoproteins.

Previous studies have indicated that Bromelain removes the Spike and haemagglutinin proteins of Semliki Forest virus, Sindbis virus, mouse gastrointestinal coronavirus, hemagglutinating encephalomyelitis virus and H1N1 influenza viruses, rendering these non-infective (13, 14). Moreover, Acetylcysteine was shown to alter the Spike protein by disrupting the three cystine disulfide bridges between Spike and Envelope proteins thereby reducing infectivity of SARS-CoV (15).

Therefore, in the current study we set out to determine whether BromAc can disrupt the integrity of SARS-CoV-2 Spike and Envelope proteins and subsequently examine its inactivation potential on *in vitro* replication of two viral strains, including a Spike mutant alteration of the novel S1/S2 cleavage site.

## Results

### Disintegration of SARS-CoV-2 Spike and Envelope proteins

Treatment of the Spike protein with Acetylcysteine alone did not show any alteration of the protein, whereas concentrations of Bromelain 50 and 100 µg/mL and BromAc 50 and 100 µg/20mg/mL resulted in protein alteration (Figure 1A). Treatment with Acetylcysteine on the Envelope protein did not alter the protein whereas treatment with Bromelain 50 and 100 µg/mL and BromAc 50 and 100 µg/20mg/mL also resulted in near complete and complete fragmentation, respectively (Figure 1A).

**Figure 1.**
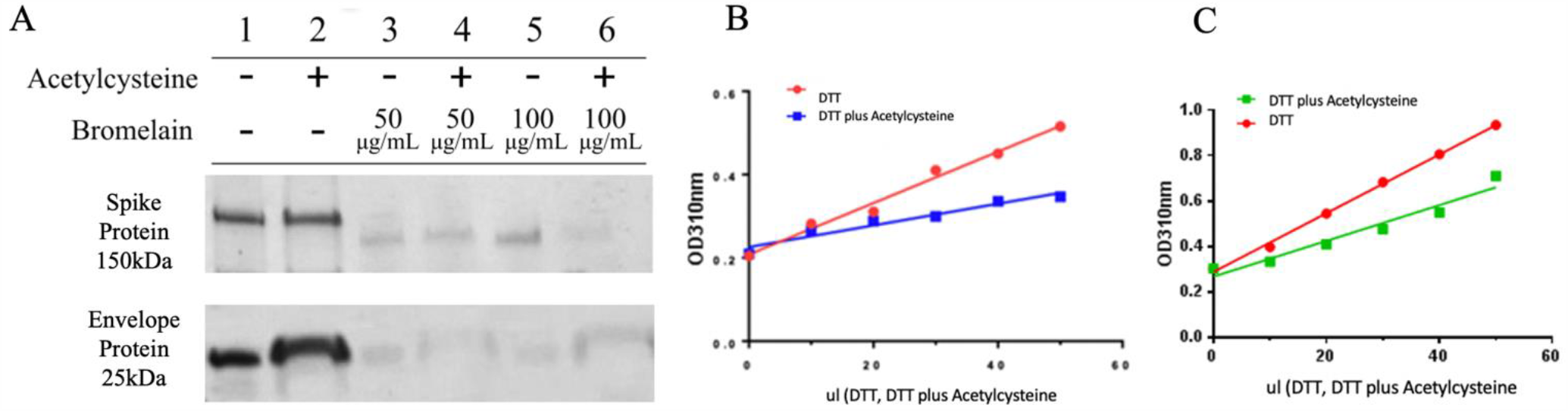
**A)** Bromelain and Acetylcysteine present a synergistic effect on SARS-CoV-2 Spike and Envelope protein destabilization. SDS-PAGE of the recombinant SARS-CoV-2 Spike protein S1+S2 subunits (150 kDa) and Envelope protein (25 kDa). Proteins were treated with 20 mg/mL Acetylcysteine alone, 100 and 50 µg/mL Bromelain alone, and combination 100 and 50 µg/20mg/mL BromAc. **B)** Disulfide reduction of recombinant SARS-CoV-2 Spike protein by Acetylcysteine. The differential assay between Acetylcysteine and DTT for the reduction of disulfide bonds found on the Spike protein indicates that Acetylcysteine reduces 42% of the disulfide bonds before the addition of DTT. The remaining bonds are reduced by DTT to produce the chromogen detected at 310 nm. **C)** Disulfide reduction of recombinant SARS-CoV-2 Envelope protein by Acetylcysteine. The differential assay between Acetylcysteine and DTT for the reduction of disulfide bonds found on the Envelope protein indicates that Acetylcysteine reduces 40% of the bonds before the addition of DTT.

### UV spectral detection demonstrates the disintegration of disulfide bonds in Spike and Envelope proteins

The comparative reduction of disulfide bonds on the Spike protein between DTT alone and DTT with Acetylcysteine demonstrated a 42% difference (Figure 1B), calculated based on the slope of the graphs [0.002599/0.006171 (100) = 42 %]. Acetylcysteine was thus able to reduce 58% of the disulfide linkages in the sample after which the remaining disulfide bonds were reduced by DTT to produce the chromogen that was monitored in the spectra. Similarly, the differential assay between Acetylcysteine and DTT for the reduction of disulfide bonds found in the Envelope protein [0.007866/0.01293 (100) = 60%] indicates that that Acetylcysteine reduces 40% of the disulfide bonds before the addition of DTT (Figure 1C).

### In vitro SARS-CoV-2 inactivating potential of Bromelain, Acetylcysteine, and BromAc

For both SARS-CoV-2 strains tested, the untreated virus controls at 10^5.5^ and 10^4.5^ TCID_50_/mL yielded typical cytopathic effects (CPE), and no cytotoxicity was observed for any of the drug combinations on Vero cells. Optical CPE results were invariably confirmed by end-point Neutral Red cell staining. Overall, Bromelain and Acetylcysteine treatment alone showed no viral inhibition, all with CPE comparable to virus control wells, whereas BromAc combinations displayed virus inactivation in a concentration-dependent manner (Figure 2). Treatment on 10^4.5^ TCID_50_/mL virus titers (Figures 2B and 2D) yielded more consistent inhibition of CPE for quadruplicates than 10^5.5^ TCID_50_/mL virus titers (Figures 2A and 2C).

**Figure 2.**
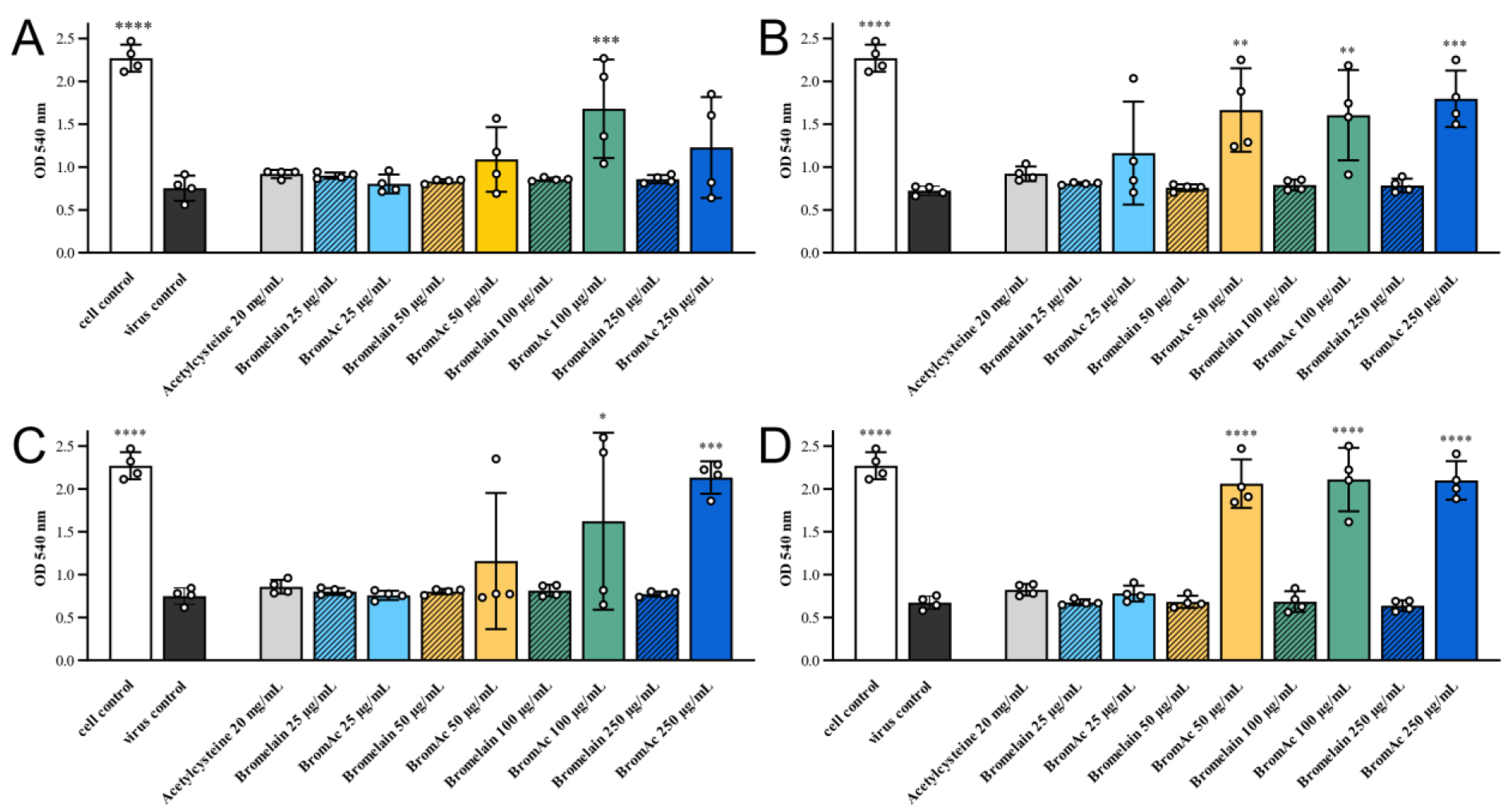
Cell lysis assays demonstrated in vitro inactivation potential of BromAc against SARS-CoV-2. Cell viability was measured by cell staining with Neutral Red, where optical density (OD) is directly proportional to viable cells. Low OD would signify important cell lysis due to virus replication. The WT SARS-CoV-2 strain at 5.5 and 4.5 log_10_TCID_50_/mL titers (A and B, respectively) showed no inhibition of cytopathic effect (CPE) for single agent treatment, comparable to the mock treatment virus control condition. BromAc combinations were able to inhibit CPE, comparable to the mock infection cell controls. Treatment of a SARS-CoV-2 S1/S2 Spike protein variant (ΔS) with a mutation at the S1/S2 junction at 5.5 and 4.5 log_10_TCID_50_/mL titers (C and D, respectively) showed similar results. Bars represent the average of the each quadruplicate per condition, illustrated by white circles. Ordinary one-way ANOVA was peformed, using the mock treatment virus control as the control condition (****p<0.0001, ***p<0.0005, **p<0.003, *p<0.05).

Based on the virus inactivation guidelines established by the World Health Organization (16), a robust and reliable process of inactivation will be able to reduce replication by at least 4 logs [Log_10_ reduction value (LRV) = (RT-PCR Ct treatment – RT-PCR Ct control)/3.3; as 1 log_10_ ≈ 3.3 Ct]. As such, RT-PCR was performed on the RNA extracts to directly measure virus replication. For the wildtype (WT) strain at 10^4.5^ TCID_50_/mL, successful LRV > 4 were observed with 1 out of 4 wells, 2 out of 4 wells, 3 out of 4 wells and 4 out of 4 wells for 25, 50, 100 and 250 µg/20mg/mL BromAc, respectively (Figure 4). It is worth noting that at 10^5.5^ TCID_50_/mL, LRV were slightly below the threshold at, on average, 3.3 with 3 out of 4 wells and 2 out of 4 wells for 100 and 250 µg/20mg/mL BromAc (Table 1). For the Spike protein mutant (ΔS) at 10^4.5^ TCID_50_/mL, no successful LRV > 4 was observed for 25 µg/20mg/mL BromAc, while it was noted in 4 out of 4 wells for 50, 100 and 250 µg/20mg/mL BromAc (Figure 4). Of note, at 10^5.5^ TCID_50_/mL, LRV were slightly below the threshold at, on average, 3.2 with 1 out of 4 wells, 2 out of 4 wells, and 4 out of 4 wells for 50, 100 and 250 µg/20mg/mL BromAc (Table 1). Overall, *in vitro* inactivation of both SARS-CoV-2 strains replication capacity was observed in a dose-dependent manner, most strongly demonstrated at 100 and 250 µg/20mg/mL BromAc against 10^4.5^ TCID_50_/mL of virus.

**Table 1.** Log_10_ reduction values (LRV) of *in vitro* virus replication 96 hours after BromAc treatment on WT and ΔS SARS-CoV-2 strains at 5.5 and 4.5 log_10_TCID_50_/mL titers. LRV were calculated with the following formula: LRV = (RT-PCR Ct of treatment – RT-PCR Ct virus control)/3.3; as 1 log_10_ ≈ 3.3 Ct. Each replicate is described. TCID_50_/mL = Median Tissue Culture Infectious Dose; WT = wild-type; ΔS = S1/S2 Spike mutant.

| | BromAc<br>($\mu\text{g}/20\text{mg}/\text{mL}$) | Virus titer | | | | | | | |
| --- | --- | --- | --- | --- | --- | --- | --- | --- | --- |
| | | 5.5 $\log_{10}\text{TCID}_{50}/\text{mL}$ | | | | 4.5 $\log_{10}\text{TCID}_{50}/\text{mL}$ | | | |
| WT | 25 | 0.033 | 0.104 | 0.250 | 0.213 | 0.463 | 0.356 | 4.390 | 0.173 |
|  | 50 | 0.050 | 0.304 | 0.446 | 0.698 | 0.471 | 4.378 | 0.404 | 4.651 |
|  | 100 | 3.415 | 3.323 | 0.360 | 3.313 | 4.418 | 4.463 | 0.423 | 4.508 |
|  | 250 | 0.033 | 3.423 | 0.200 | 3.389 | 4.496 | 4.370 | 4.419 | 4.506 |
| $\Delta\text{S}$ | 25 | 0.010 | 0.153 | NA | 0.414 | 0.330 | 0.313 | 0.172 | 0.075 |
|  | 50 | 3.252 | 0.297 | 0.278 | 0.275 | 4.762 | 4.612 | 4.618 | 4.571 |
|  | 100 | 3.191 | 3.260 | 0.210 | 0.301 | 6.054 | 4.518 | 5.155 | 4.747 |
|  | 250 | 3.287 | 3.298 | 3.308 | 3.308 | 4.333 | 4.302 | 4.410 | 4.361 |

**Figure 3.**
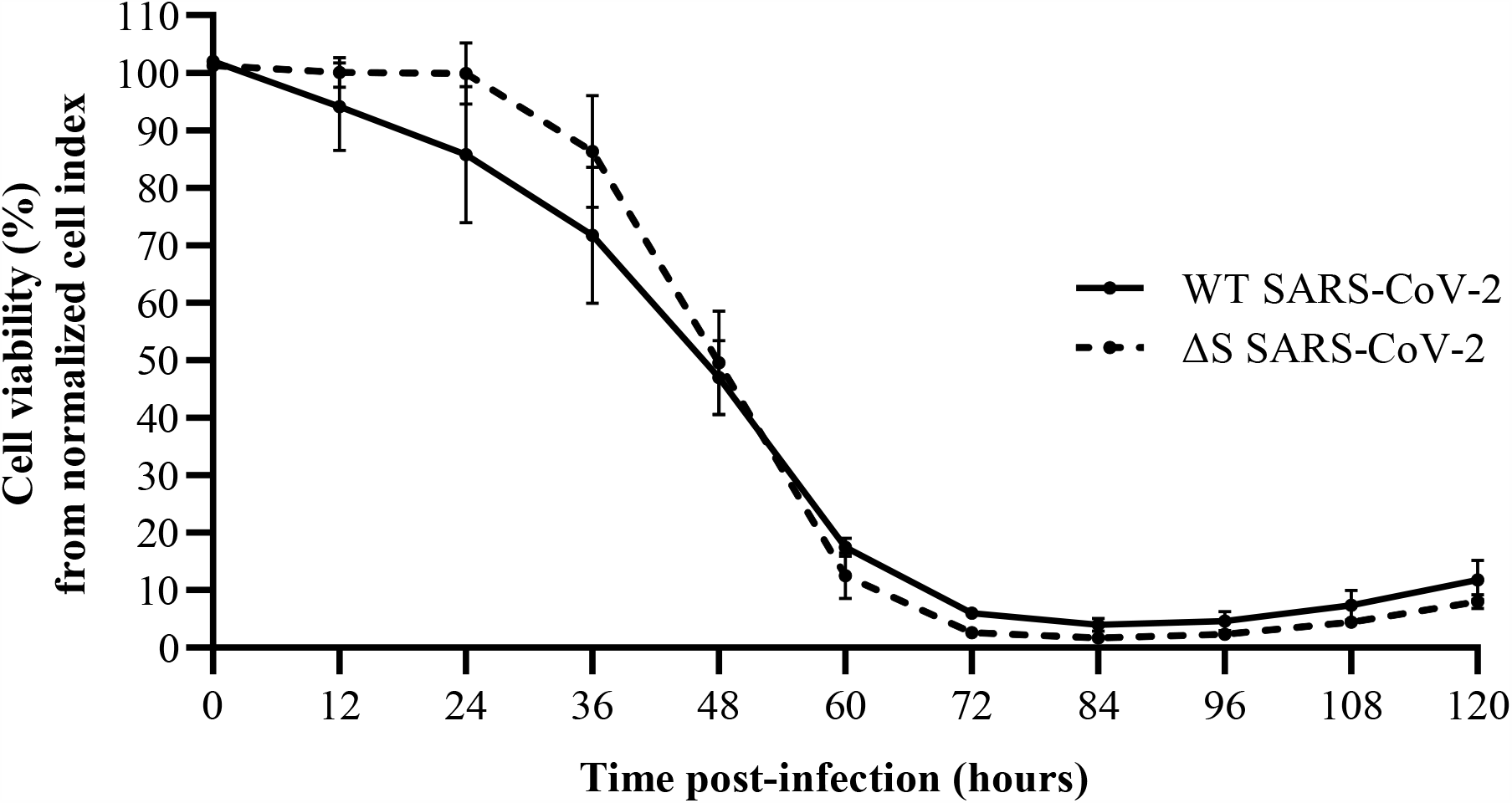
SARS-CoV-2 replication capacity of WT and ΔS SARS-CoV-2 by Real-Time Cell Analysis. Data points correspond to area under the curve analysis of normalized cell index (electronic impedance of RTCA established at time of inoculation) at 12-hour intervals. Then, cell viability was determined by normalizing against the corresponding cell control. WT = wild-type; ΔS = S1/S2 Spike mutant.

**Figure 4.**
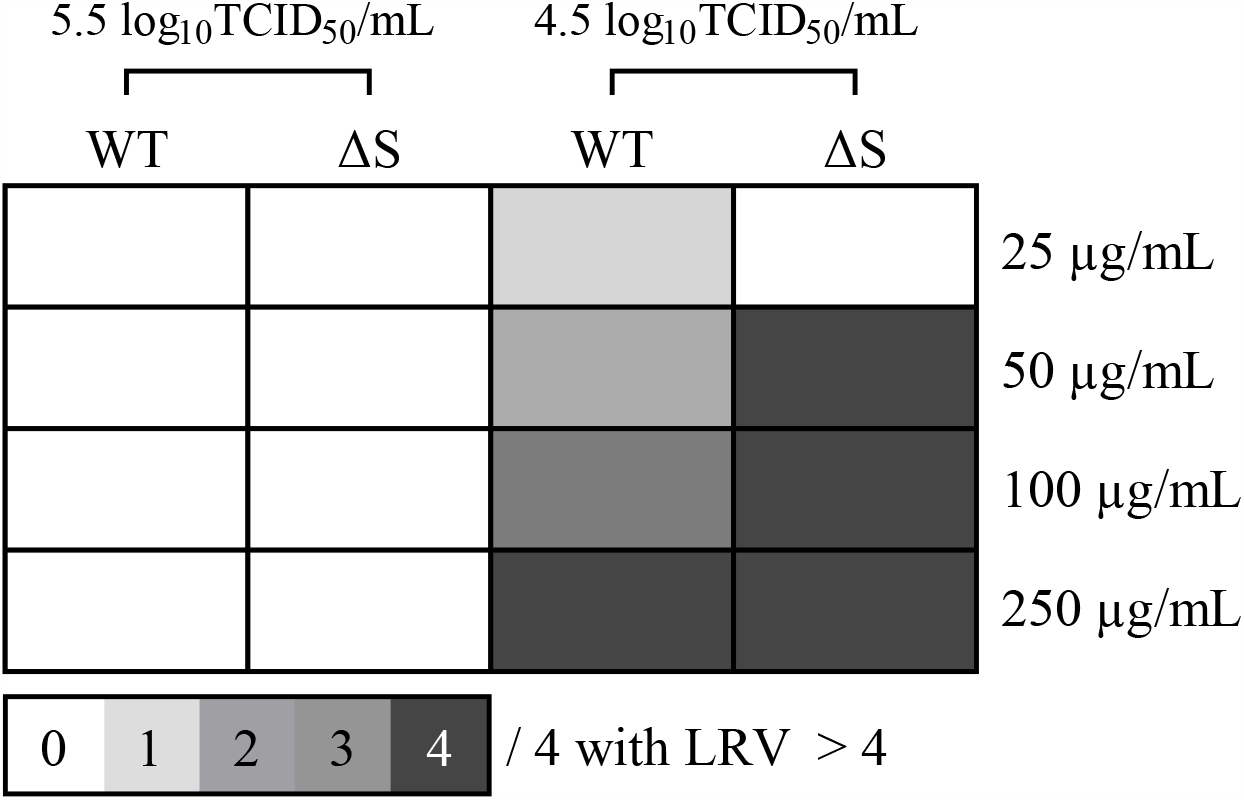
Threshold matrix of log_10_ reduction values (LRV) of *in vitro* virus replication 96 hours after BromAc treatment on WT and ΔS SARS-CoV-2 strains at 5.5 and 4.5 log_10_TCID_50_/mL titers. LRV were calculated with the following formula: LRV = (RT-PCR Ct of treatment – RT-PCR Ct virus control)/3.3; as 1 log10 ≈ 3.3 Ct. The color gradient matrix displays the number of quadruplicates per condition yielding an LRV>4, corresponding to a robust inactivation according to the WHO. WT = wild-type; ΔS = S1/S2 Spike mutant.

Real-time cell analysis demonstrated comparable replication kinetics for both WT and ΔS SARS-CoV-2 strains (Figure 3). No significant difference of cell viability was observed between WT and ΔS at any time point. From 48 hours post-infection, WT and ΔS cell viability was significantly different to the mock infection (p<0.05).

## Discussion

The combination of Bromelain and Acetylcysteine, BromAc, synergistically inhibited the infectivity of two SARS-CoV-2 strains cultured on Vero cells. A hypothesized mechanism of action could be the disruption of the Spike glycoprotein and the reduction of its disulfide bonds.

The SARS-CoV-2 Spike protein is the cornerstone of virion binding to host cells and hence represents an ideal therapeutic target. A direct mechanical action against this Spike protein is a different treatment strategy in comparison to most of the existing antiviral drugs: preventing viral entry in host cells rather than targeting the replication machinery. BromAc acts as a biochemical agent to destroy complex glycoproteins. Bromelain’s multipotent enzymatic competencies, dominated by the ability to disrupt glycosidic linkages, usefully complete Acetylcysteine’s strong power to reduce disulfide bonds (9). Amino acid sequence analysis of the SARS-CoV-2 Spike glycoprotein identified several predetermined sites where BromAc could preferentially act, such as the S2’ site rich in disulfide bonds (17), together with three other disulfide bonds in the receptor binding domain (RBD) (18). In parallel, the role of the glycosidic shield in covering the Spike, prone to be removed by BromAc, has been highlighted as a stabilizer of the molecular structure and a resistance mechanism to specific immune response (5, 19).

Mammalian cells exhibit reductive functions at their surface capable of cleaving disulfide bonds, and the regulation of this thiol-disulfide balance has been proven to impact the internalization of different type of viruses (20-23). This internalization process is the key mechanism of enveloped virus infection and is based on dynamic conformational changes of their surface glycoproteins, namely mediated by disulfide bond reduction, regulated by cell surface oxydoreductases and proteases (5, 24-27). Two distinct conformations of the Spike, more specifically of its RBD, are possible. The “down” conformation is required for ACE2 binding to RBD (5). Both ACE2 and Spike proteins possess disulfide bonds (24). The binding induces Spike proteolysis by TMPRSS2 or Cathepsin L triggering the conformational modifications to the post-fusion state (5). The energy liberated by the disulfide bond reduction increases protein flexibility, which is maximal when the reduced status is complete (24), and thus allowing the fusion of host-virus membranes, otherwise impossible due to the repulsive hydration forces present before reduction(5). Conversely, Hati and Bhattacharyya demonstrated that when all the RBD disulfide bonds were reduced to thiols, the receptor binding became less favorable (24). Interestingly, the reduction of ACE2 disulfide bonds also induced a decrease of binding (24). Moreover, other reports suggested that Bromelain alone could inhibit SARS-CoV-2 infection in VeroE6 cells through an action on disulfide links (28). As such, the loss of SARS-CoV-2 infectivity observed after pre-treatment by BromAc could be correlated to the cumulative destruction of the Spike and Envelope proteins, with a significant reduction of their disulfide bonds by Acetylcysteine, demonstrated *in vitro*.

Interestingly, a similar effect of BromAc was observed against both WT and ΔS SARS-CoV-2. The main difference in amino acid sequences between SARS-CoV-2 and previous SARS-CoV is the inclusion of a furin cleavage site between S1 and S2 domains (29). This distinct site of the Spike protein and its role in host spill-over and virus fitness is a topic of much debate (29-32). Of note, ΔS, which harbors a mutation in this novel S1/S2 cleavage site and alters the cleavage motif, was also sensitive to BromAc treatment, with no observed difference in replication capacity to the WT. These results would suggest that, from a threshold dose, BromAc could potentially be effective on Spike mutant strains. This may be a clear advantage for BromAc over specific immunologic mechanisms of a Spike-specific vaccination (3, 4).

To date, different treatment strategies have been tested, but no molecules have demonstrated a clear antiviral effect. In addition, given the heterogeneous disease outcome of COVID-19 patients, the treatment strategy should combine several mechanisms of action and be adapted to the stage of the disease. Thus, treatment repurposing remains an ideal strategy against COVID-19, whilst waiting for sufficient vaccination coverage worldwide (33, 34). In particular, the development of early nasal-directed treatment prone to decrease a patient’s infectivity and prevent the progression towards severe pulmonary forms is supported by a strong rationale. Hou et al demonstrated that the first site of infection is nasopharyngeal mucosa with a secondary movement to the lungs by aspiration (35). Indeed, the pattern of infectivity of respiratory tract cells followed ACE2 receptor expression, decreasing from the upper respiratory tract to the alveolar tissue. The ratio for ACE2 was 5-fold greater in the nose than the distal respiratory tract (36). Other repurposing treatments as a nasal antiseptic have been tested *in vitro*, such as Povidone-Iodine, which has shown activity against SARS-CoV-2 (37). In the present study, we showed the *in vitro* therapeutic potential of BromAc against SARS-CoV-2 with a threshold efficient dose at 100 µg/20mgmL BromAc. As animal airway safety models in two species to date have exhibited no toxicity (*Unpublished data*), the aim is to test nasal administration of the drug in a phase I clinical trial (ACTRN12620000788976). Such treatment could help mitigate mild infections and prevent infection of persons regularly in contact with the virus, such as health-care workers.

Although our results are encouraging, there are a number of points to weigh up this demonstration. Namely, the *in vitro* conditions are fixed and could be different from *in vivo*. Any enzymatic reaction is influenced by the pH of the environment, all the more when it concerns redox reactions such as disulfide bond reduction (25). The nasal mucosal pH is physiologically comprised between 5.5 and 6.5 and increases in rhinitis to 7.2-8.3 (38). Advanced age, often encountered in SARS-CoV-2 symptomatic infections, also induces a nasal mucosa pH increase (38). Such a range of variation, depending on modifications typically induced by a viral infection, may challenge the efficacy of our treatment strategy. Further *in vitro* experiments to test various conditions of pH are ongoing, but ultimately, only clinical studies will be able to assess this point. The type of cells targeted by BromAc is also of importance. Our experiments were led on a monkey kidney cell line known to mimic main factors influencing SARS-CoV-2 infectivity. With the above hypotheses of S protein lysis thiol-disulfide balance disruption, BromAc efficacy should not be influenced by the membrane protease pattern, as is the case for Chloroquine (39). Reproducing this experimental protocol with the human pulmonary epithelial Calu-3 cell line would allow these points to be addressed, as virus entry is TMPRSS2-dependent and pH-independent, as in airway epithelium, while virus entry in Vero cells is Cathepsin L dependent, and so pH-dependent (39).

Overall, results obtained from the present study in conjunction with complimentary studies on BromAc properties and SARS-CoV-2 characterization reveal a strong indication that BromAc can be developed into an effective therapeutic agent against SARS-CoV-2.

## Materials and Methods

### Materials

Bromelain API was manufactured by Mucpharm Pty Ltd (Australia) as a sterile powder. Acetylcysteine was purchased from Link Pharma (Australia). The recombinant SARS-COV-2 Spike protein was obtained from SinoBiological (Cat#40589-V08B1). The recombinant Envelope protein was obtained from MyBioSource (Cat#MBS8309649). All other reagents were from Sigma Aldrich.

### Recombinant Spike and Envelope gel electrophoresis

The Spike or Envelope protein was reconstituted in sterile distilled water according to the manufacturer’s instructions and aliquots were frozen at −20^0^C. Two and a half micrograms of Spike or Envelope protein were incubated with 50 or 100 µg/mL Bromelain, 20 mg/mL Acetylcysteine or combination in Milli-Q water. Control contained no drugs. The total reaction volume was 15 µL each. After 30 mins incubation at 37^0^C, 5 µl of sample buffer was added into each reaction. A total of 20 µl of each reaction was electrophoresed on a SDS-PAGE (Bio-Rad; Cat#456-1095). The gels were stained using Coomassie blue.

### UV Spectral detection of disulfide bonds in Spike and Envelope proteins

The recombinant SARS-CoV-2 Spike protein at a concentration of 3.0 µg/mL in PBS (pH 7.0) containing 1 mM EDTA was incubated with 0, 10, 20, 40, and 50 µL of Acetylcysteine (0.5 M) and agitated at 37°C for 30 min followed by equivalent addition of DTT (Dithiothreitol) (0.5 M) and agitated for a further 30 min at 37°C. The Spike protein was in parallel incubated only with DTT (0.5 M) as before without any Acetylcysteine and agitated at 37°C for 30 min. The absorbance was then read at 310 nm. UV spectral detection of disulfide bonds in the Envelope protein was performed in a similar manner.

### SARS-CoV-2 whole virus inactivation with BromAc

Fully respecting the WHO interim biosafety guidance related to the coronavirus disease (40), the SARS-CoV-2 whole virus inactivation tests were carried out with a wild type (WT) strain representative of early circulating European viruses (GISAID accession number EPI_ISL_578176). A second SARS-CoV-2 strain (denoted as ΔS), reported through routine genomic surveillance in the Auvergne-Rhône-Alpes region in France, was added to the inactivation tests due to a rare mutation in the Spike S1/S2 cleavage site and its culture availability at the laboratory (GISAID accession number EPI_ISL_578177).

These tests were conducted with incremental concentrations of Bromelain alone (0, 25, 50, 100 and 250 µg/mL), Acetylcysteine alone (20 mg/mL), and with the cross-reaction of the different Bromelain concentrations combined to a constant 20 mg/mL Acetylcysteine formulation, against 2 virus culture dilutions at 10^5.5^ and 10^4.5^ TCID_50_/mL. Following 1 hour of drug exposure at 37°C, all conditions, including the control, were diluted 100-fold to avoid cytotoxicity, inoculated in quadruplicate on confluent Vero cells (ATCC©, CCL-81), and incubated for 5 days at 36°C with 5% CO_2_. Cells were maintained in Eagle’s minimal essential medium (EMEM) with 2% Penicillin-Streptomycin, 1% L-glutamine, and 2% inactivated fetal bovine serum. Results were obtained by daily optical microscopy observations, an end-point cell lysis staining assay, and reverse-transcriptase polymerase chain reaction (RT-PCR) of supernatant RNA extracts. Briefly, the end-point cell lysis staining assay consisted of adding Neutral Red dye (Merck KGaA, Darmstadt, DE) to cell monolayers, incubating at 37°C for 45 minutes, washing with PBS, and adding citrate ethanol before optical density (OD) was measured at 540 nm (Labsystems Multiskan Ascent Reader, Thermo Fisher Scientific). OD was directly proportional to viable cells, so a low OD would signify important cell lysis due to virus replication. In addition, RNA from well supernatants was extracted by the semi-automated eMAG® workstation (bioMérieux, Lyon, FR), and SARS-CoV-2 RdRp IP2-targeted RdRp Institute Pasteur RT-PCR was performed on a QuantStudio™ 5 System (Applied Biosystems, Thermo Fisher Scientific). Log_10_ reduction values (LRV) of viral replication were calculated by the difference between treatment and control wells per condition divided by 3.3 (as 1 log_10_ ≈ 3.3 PCR Cycle thresholds (Ct)).

### Replication Kinetics by Real-Time Cell Analysis

To compare the *in vitro* replication capacity of both WT and ΔS SARS-CoV-2 strains, replication kinetics was determined by measuring electrode impedance of microelectronic cell sensors on the xCELLigence Real-Time Cell Analyzer (RTCA) DP Instrument (ACEA Biosciences, Inc.). Vero cells (ATCC^©^, CCL-81) were seeded at 20,000 cells per well on an E-Plate 16 (ACEA Biosciences, Inc., CA, USA) and incubated with the same media conditions as described previously at 36°C with 5% CO_2_. After 24 hours, SARS-CoV-2 culture isolates were inoculated in triplicate at a multiplicity of infection of 10^−2^. Mock infections were performed in quadruplicate. Electronic impedance data (cell index) was continuously collected at 15-minute intervals for 6 days. Area under the curve analysis of normalized cell index, established at time of inoculation, was then calculated for 12-hour intervals. At each interval, cell viability was determined by normalizing against the corresponding cell control. Tukey multiple comparison tests were used to compare each condition on GraphPad Prism (software version 9.0).

